# Range Overlap Drives Chromosome Inversion Fixation in Passerine Birds

**DOI:** 10.1101/053371

**Authors:** Daniel M. Hooper

## Abstract

Chromosome inversions evolve frequently but the reasons why remain largely enigmatic. I used cytological descriptions of 410 species of passerine birds (order Passeriformes) to identify pericentric inversion differences between species. Using a new fossil-calibrated phylogeny I examine the phylogenetic, demographic, and genomic context in which these inversions have evolved. The number of inversion differences between closely related species was highly variable yet consistently predicted by a single factor: whether the ranges of species overlapped. This observation holds even when the analysis is restricted to sympatric sister pairs known to hybridize, and which have divergence times estimated similar to allopatric pairs. Inversions were significantly more likely to have fixed on a sex chromosome than an autosome yet variable mutagenic input alone (by chromosome size, map length, GC content, or repeat density) cannot explain the differences between chromosomes in the number of inversions fixed. Together, these results support a model in which inversions in passerines are adaptive and spread by selection when gene flow occurs before reproductive isolation is complete.

## Introduction

Speciation occurs not just from the accumulation of molecular changes in DNA composition but also with physical rearrangements to the structure of diverging genomes. Chromosome inversions, one common type of chromosomal rearrangement, are often observed as fixed differences between species and as polymorphisms segregating within species [1,2]. Chromosome inversions are thought to impact sex chromosome evolution [3–5], supergene formation [6–9], local adaptation [10–13], and reproductive isolation [14–18]. Despite their potential evolutionary importance, the widespread presence of inversions is puzzling, as new rearrangements may be initially disfavored due to structural underdominance in heterokaryotypes, if crossing over within the inverted region during meiosis results in the production of aneuploid gametes [1,2,19]. Reconciling the pervasiveness of chromosome inversions both between and within species with possible selective disadvantages that a new inversion faces remains an unresolved problem.

Traditional models of inversion fixation rely upon genetic drift acting within highly structured populations to lift an inversion above 50% frequency in a deme, after which selection favors its spread [20–24]. This model predicts that inversion fixation should be independent of population size [20]. Hooper and Price [25] tested this in the Estrildid finches (family Estrildidae) and rejected it, based on the strong positive relationship observed between the rate of inversion fixation and range size. Other models of inversion fixation have focused on selection, in which drift plays no part. An inversion may be adaptive and spread (1) if its breakpoints favorably alter gene expression [26,27] or (2) by meiotic drive if it happens to link alleles that together alter segregation distortion [28,29]. Alternatively, recent models emphasize the contextual selective advantage of an inversion if it suppresses recombination [30–32]. In these scenarios, an inversion will spread when natural selection favors the maintenance of linkage disequilibrium between sets of alleles that are locally adapted to (3A) the habitat or (3B) the genetic background of a population that would otherwise be broken up by recombination.

The alternative selection models make different predictions that can be tested with comparative analyses. All adaptive models depend on mutational input, which should scale with population size, but selection pressures arising from breakpoint effects and meiotic drive models should be particularly strongly associated with mutational input. This is because selection pressures are less dependent on environmental context and mutations that produce favorable effects are likely to be rare. Hence, these models predict a strong scaling with population size. On the other hand, in recombination suppression models gene flow plays a central role, as it generates the selective advantage for a new inversion. In one case (hypothesis 3A, above), ecological differences are essential if local adaptation is to create a selective advantage for an inversion [31]. Ecological differences can be assessed by measuring characteristics such as body mass, feeding guild, habitat associations, etc. Gene flow in the form of hybridization between incompletely reproductively isolated forms is required if inversions are to increase in response to genetic incompatibilities (hypothesis 3B). This latter model therefore predicts partially reproductively isolated forms should have had the potential to interbreed.

In Estrildidae, a previous analysis showed that both range size and range overlap were positively associated with the rate of inversion fixation [25]. However, range size and range overlap were themselves correlated, and it was difficult to disentangle their contributions, given the sample size (N = 32 species). The range size of a species may be considered an estimate of population size, predicted to be a strong correlate in the breakpoint and meiotic drive models. Range overlap between closely related species indicates the potential for hybridization, predicted to be essential in genetic incompatibility models. The distinction between the two range effects is important as it suggests alternative adaptive roles for chromosome inversions: range overlap implies that inversion evolution may be intimately linked to the speciation process through interspecific interactions (3B), but without range overlap this is not possible.

The Passeriformes are just one of 39 extant orders of birds yet comprise over half of all avian biodiversity with the ~6000 species found in nearly every terrestrial habitat on the planet [33]. The rapid rate of speciation and geographic expansion in passerines is coupled with extensive eco-morphological diversification: body size varies >350-fold between the smallest and largest species (4.2 g to 1,500 g) while variation in beak shape and behavior has produced a wide spectrum of feeding morphologies (nectarivores, granivores, insectivores, frugivores, etc. [33,34]). In contrast to the wealth of eco-morphological diversity in passerines, the gross structure of the passerine genome does not vary greatly, with diploid chromosome number (2N) falling between 76–80 for 77% of species (Table S1; reviewed in [35]). Although inter-chromosomal rearrangements like chromosome fusions, fissions, and translocations may be generally rare in birds, inversions are far more common [25,35–38]. Cytological evidence for the frequent occurrence of inversions in birds is corroborated by recent genomic studies that observe hundreds of inversion-derived rearrangements between species [39–45].

In order to evaluate support for alternative models of inversion fixation, I here use the phylogenetic and genomic distribution of large pericentric inversions (i.e. those encompassing the centromere) identified from the cytological literature of 410 species of passerine birds. To this end, I build a fossil-calibrated phylogeny of those passerines with karyotype data and map pericentric inversion fixation on the 9 largest autosomes and both sex chromosomes (S1-4 Tables). I compare the extent of inversion differentiation, across 80 passerine clades and 47 sister species pairs, with differences between species in range size and their patterns of range overlap (S5-7 Tables). I evaluate the importance of raw mutagenic input on the genomic distribution of inversions by assessing how the number of inversions on a chromosome is associated with its size, map length, repeat density, and GC content or whether it was an autosome or sex chromosome (S8-9 Tables). By examining the drivers of chromosome inversion evolution in a group as species rich and ecologically diverse as the passerines, I aim to illuminate both the frequency with which inversions occur in birds and the implications for chromosome inversions and speciation with gene flow, broadly.

## Results

### Phylogenetic Signal of Chromosome Inversion Evolution in Passerines

The time-calibrated phylogeny for the 410 karyotyped passerine species in this study is shown in Figure 1 with family memberships labeled (see also S1-2 Figs.). The topology is congruent both between and within families to previously published studies in Passeriformes (S5 Table). Divergence time estimates between families are consistent again with recent phylogenomic studies of crown Aves that utilized partially overlapping fossil sets and similar calibration methods [46,47].

**Fig 1.**
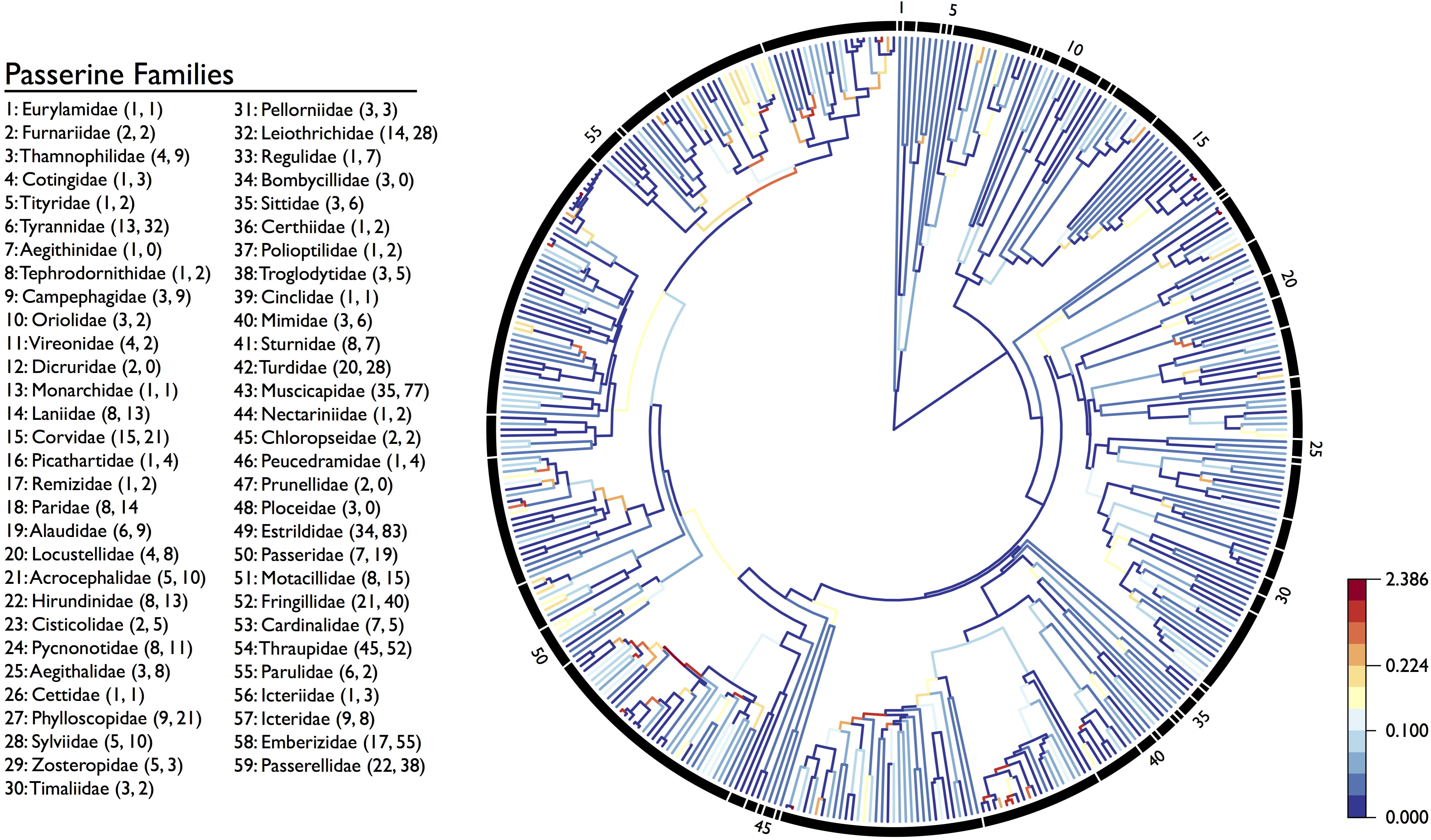
Pericentric inversion fixation rate variation across Passeriformes. Passerine families included in this study are shown on the left: numbers within parentheses refer to the number of karyotyped species and number of pericentric inversions within each family, respectively. Families are ordered clockwise by phylogenetic position in the tree. The time-dated phylogeny for the 410 karyotyped species used in this study is shown on the right. Branches are color-coded according to the inferred rate of pericentric inversion fixation using the R package ggtree [48] with rates partitioned according to the Jenks natural breaks method where variance within bins is minimized, while variance between bins is maximized [49].

The rate of inversion fixation averaged across all 410 passerine species was one inversion every 5.3 million years of evolution along a branch (4269My total branch length/808 inferred inversions). Inversion fixation rate varied greatly across lineages (Fig. 1). Rates ranged from no inversions fixed over 23.7My on the lineage leading to the common iora (*Aegithina tiphia*) to an inversion on the 6^th^ largest autosome that separated the pied and black-eared wheatear (*Oenanthe pleschanka* and *O. melanoleuca*) with a divergence time of ~0.2Ma. Explaining the underlying evolutionary basis of this rate variation was the guiding motivation of this study.

### Inversion Differentiation Across 80 Passerine Clades

Analysis of inversion fixation using 80 independent passerine clades strongly suggests that time and range overlap – rather than range size – are the best predictors of pericentric inversion evolution in Passeriformes. The best model to predict the number of inversions fixed in each clade contained two variables (branch length and range overlap; S9 Table). The two top models had similar AICc scores and model weights (ΔAICc <2; S9 Table) so I used model averaging to combine them into a final model (Table 1). This averaged model included median clade range size as an additional parameter, however only branch length (p < 0.0001) and range overlap were significant (p < 0.0001; Table 1). Older clades with more sympatric species have significantly more inversions than younger and more allopatric clades (Fig. 2; Table 1) and neither range size nor division of clades by ecological niche contributed (S9 Table) Results were consistent regardless of whether I used a more relaxed ΔAICc cutoff to model averaging (i.e. averaging all models with ΔAICc <4) or an alternative minimum range overlap cutoff values (10% or 15% pairwise range overlap) to calculate the extent of range overlap in each clade.

**Table 1.**
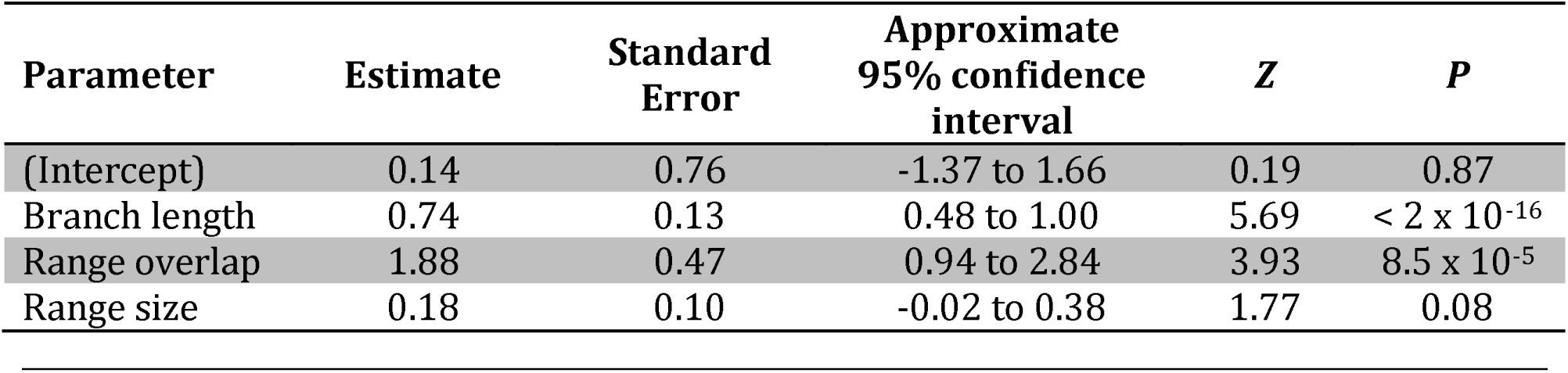
Final averaged model for clade level analysis. Phylogenetic least squares model to predict the number of pericentric inversions fixed in 80 passerine clades. Approximate 95% confidence intervals were calculated as the parameter estimate +/− 2 × standard error. P values for parameter significance were calculated 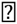 by MuMIn in R [50] model averaging the top two models with ΔAICc <2.

| Parameter | Estimate | Standard Error | Approximate 95% confidence interval | Z | P |
| --- | --- | --- | --- | --- | --- |
| (Intercept) | 0.14 | 0.76 | -1.37 to 1.66 | 0.19 | 0.87 |
| Branch length | 0.74 | 0.13 | 0.48 to 1.00 | 5.69 | $< 2 \times 10^{-16}$ |
| Range overlap | 1.88 | 0.47 | 0.94 to 2.84 | 3.93 | $8.5 \times 10^{-5}$ |
| Range size | 0.18 | 0.10 | -0.02 to 0.38 | 1.77 | 0.08 |

**Fig 2.**
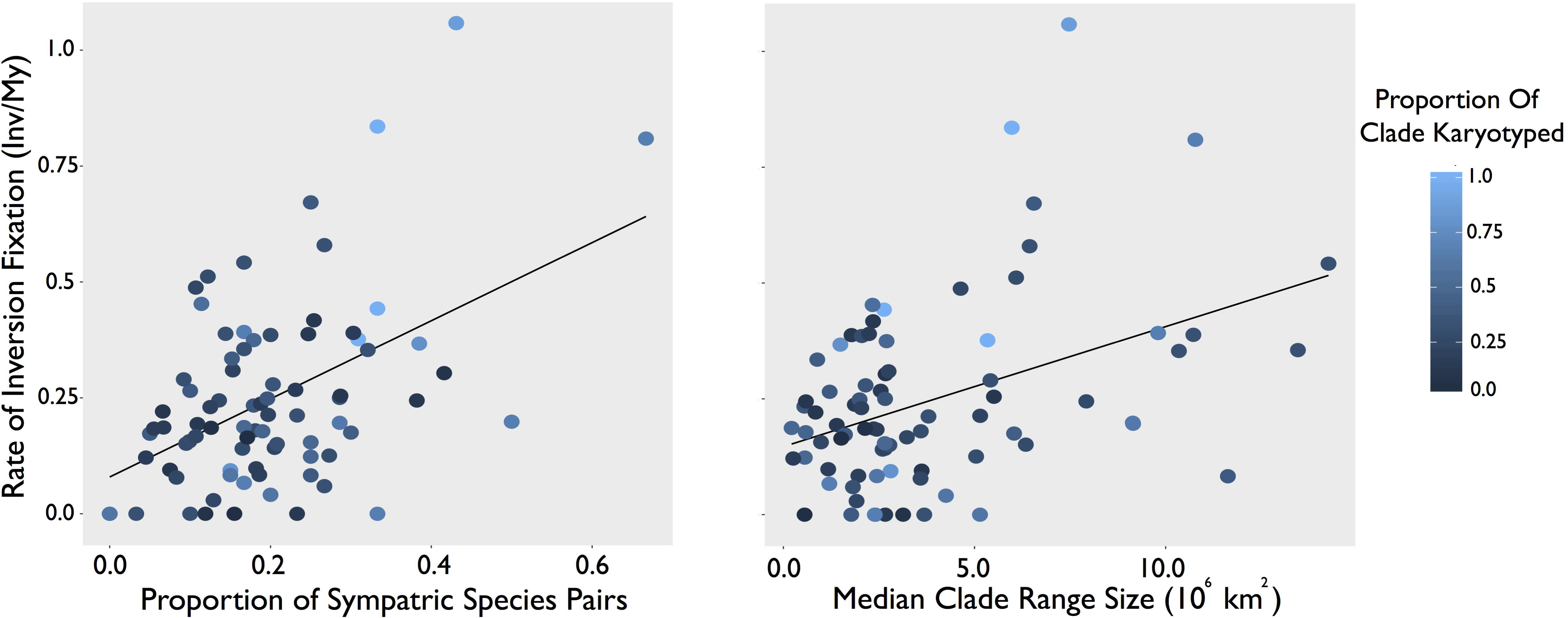
Pericentric inversion fixation rate variation across 80 passerine clades. Fixation rate is shown against the proportion of sympatric species pairs (left) and median clade range size (right). Fixation rate calculated as the total number of inversions on all chromosomes divided by the total clade branch length summed across each chromosome. Each clade is represented by a circle and shaded according to the proportion of total species with karyotype data.

### Inversion Differentiation Across 47 Sister Species Pairs

As a complement to the clade level analysis, I considered sister pairs, as clearly independent points. Sympatric sister species were significantly more likely to differ by an inversion than allopatric sisters (two-tailed *t*-test: *t*_45_ = 3.1, p = 0.003; Fig. 3A). The best model to explain the number of inversion differences between sister species contained a single parameter: whether sister species overlapped in range or not (Fig. 3B; S9 Table). The two top models had similar AICc scores and model weights (ΔAICc <2; S9 Table) so I used model averaging to combine them into a final model (Table 2). While this averaged model included sister pair age as an additional parameter, only range overlap was found to be significant (p = 0.01; Tables Table 2, 3). The 12 sister species known to hybridize in nature were more likely to be differentiated by an inversion than their allopatric counterparts (two-tailed *t*-test: *t*_21_ = 3.0, p = 0.007; Fig. 3A) despite not being genetically more divergent (*t*_21_ = 0.28, p > 0.1; S6 Table).

**Table 2.** Final averaged model for sister species analysis. Generalized linear model with Poisson errors to predict the number of pericentric inversion differences between 47 sister species pairs. Approximate 95% confidence intervals were calculated as the parameter estimate +/ − 2 × standard error. P values for parameter significance were calculated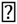 by MuMIn in R [50] after model averaging the top two models with ΔAICc <2.

| Parameter | Estimate | Standard Error | Approximate 95% confidence interval | Z | P |
| --- | --- | --- | --- | --- | --- |
| (Intercept) | -0.54 | 0.23 | -1.0 to -0.08 | 2.31 | 0.021 |
| Range overlap (Yes) | 0.67 | 0.25 | 0.17 to 1.17 | 2.57 | 0.01 |
| Sister age | 0.08 | 0.10 | -0.12 to 0.28 | 0.77 | 0.44 |

**Table 3.** Pericentric inversion differentiation between sister species. Species pairs binned by range overlap (Allopatric versus Sympatric) and inversion presence (no inversions or some inversions). The average value and range is shown for Age, Range Size, and number of Inversion Differences. Both fixed inversion differences and inversion polymorphisms observed in one sister but not the other were included. Full data in S5 Table.

| Sister Species Distribution | Inversion Presence | Sister Pairs (N) | Age (Ma) | Range Size ($10^6$ km<sup>2</sup>) | Inversion Differences |
| --- | --- | --- | --- | --- | --- |
| Allopatric | None | 6 | 2.4 (0.9 – 5.0) | 2.5 (0.2 – 5.3) | 0 |
|  | Some | 3 | 2.8 (1.0 – 6.9) | 4.0 (1.1 – 5.6) | 2.3 (1 – 4) |
| Sympatric | None | 7 | 4.25 (2.3 – 7.5) | 6.8 (2.2 – 10.9) | 0 |
|  | Some | 31 | 3.7 (0.2 – 9.3) | 7.9 (0.8 – 35.2) | 3.2 (1 – 12) |

**Fig 3.**
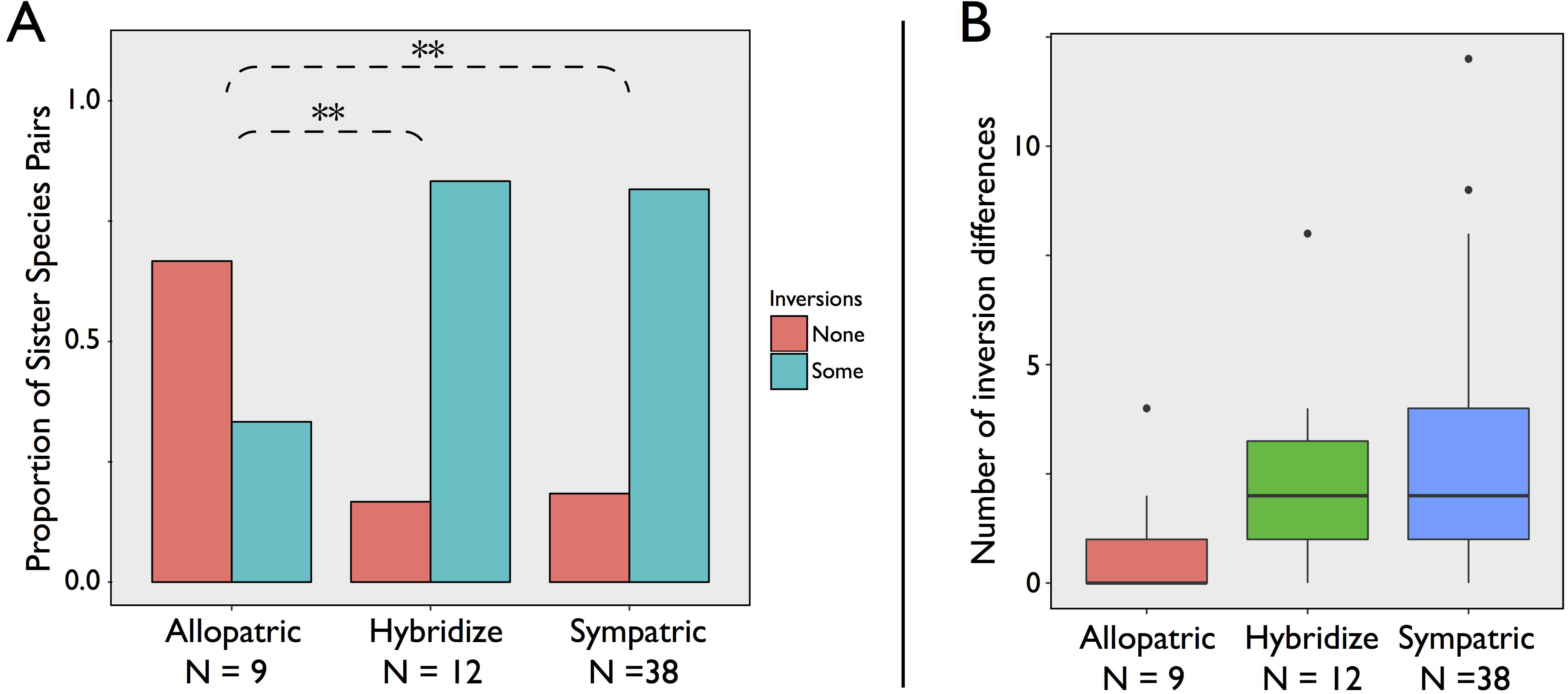
Pericentric inversion differentiation between sister species. Sister species sorted into allopatric pairs, sympatric pairs (any amount of range overlap), and the subset of sympatric sister pairs that are known to hybridize [51]. Variation between sister species groups in A) the proportion of sister pairs with and without inversion differences (none or some, respectively) and B) the number of inversion differences between sister species.

Species triplets consist of a sister pair and an outgroup species, where the outgroup overlaps one of the two sisters but not the other. Comparisons of differences between the outgroup and each sister therefore test for a role of sympatry, with time completely factored out [52–54]. Figure 4 shows a representative example where the extent of inversion differentiation is conditional upon geographic overlap with the outgroup. Results form triplet comparison confirmed the importance of range overlap (S7 Table). A conservative triplet set, where the outgroup shows no overlap with one of the sisters, has little power (N = 5) but the three triplets that show differences in the extent of inversion accumulation all find that the sister species whose range overlapped with the outgroup had accumulated more inversion differences. In a relaxed triplet set, in which some degree of range overlap was allowed between the outgroup or sisters (N = 19), 7 triplets showed no difference in inversion differentiation. However, ten triplets showed more inversions in the sister species that overlaps with the outgroup and 2 triplets the opposite pattern (two-tailed sign rank test, p = 0.039). Differences in range size between sisters did not predict the number of inversion differences (regression of contrasts, forced through the origin, p > 0.1).

**Fig 4.**
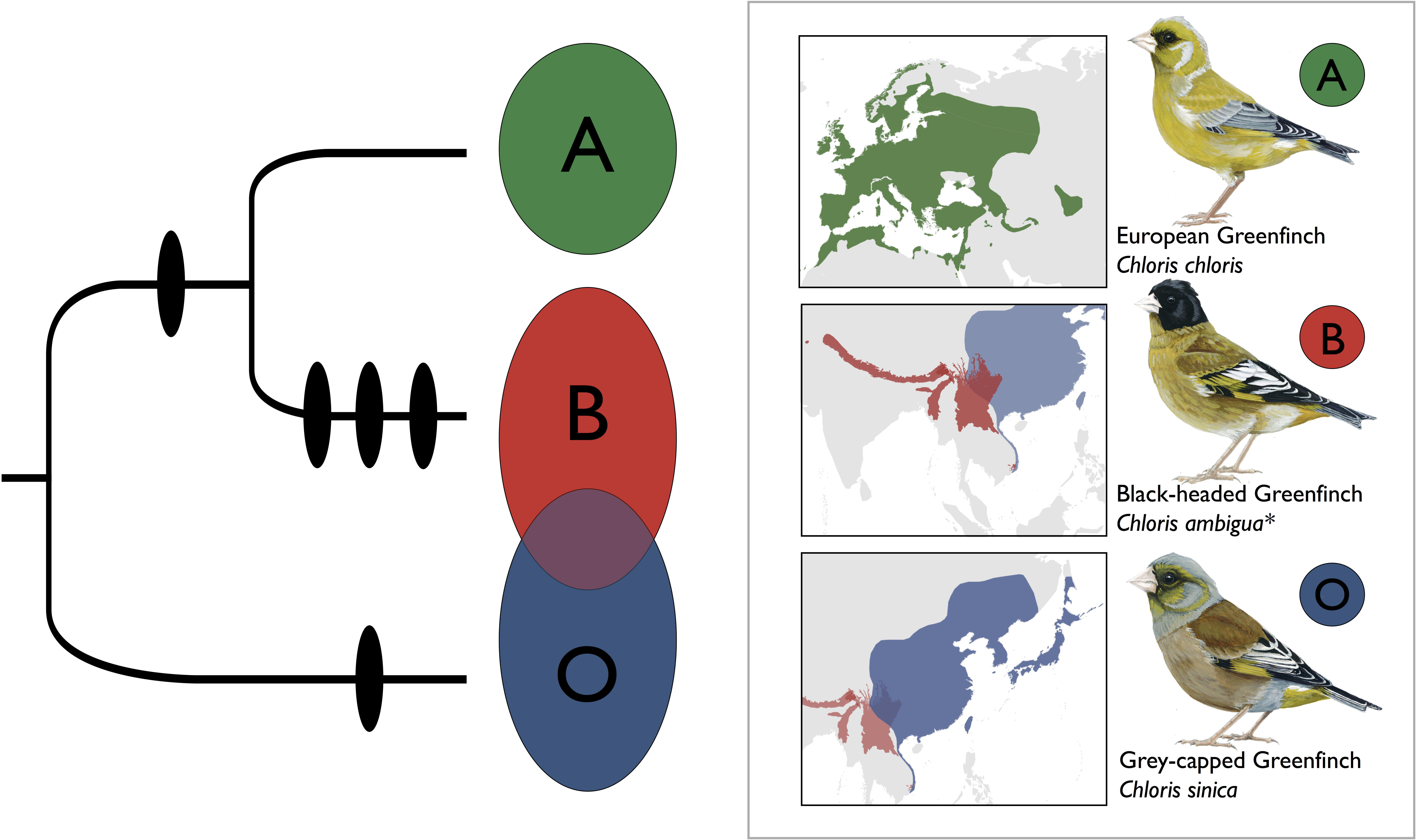
Triplet analysis of pericentric inversion evolution across greenfinches in the genus *Chloris*. The phylogenetic history of pericentric inversion fixation in *Chloris* is shown on the left, with inversions (black ovals) on the branches they are inferred to have fixed. The geographic distribution and likeness of each species is shown on the right. The European greenfinch *Chloris chloris* (green, A) and black-headed greenfinch *C. ambigua* (red, B) are allopatric sister species. The grey-capped greenfinch *C. sinica* (blue, Out) is the outgroup geographically isolated from *C. chloris* but in geographic contact with *C. ambigua*. While 5 inversions have evolved to differentiate *C. ambigua* from *C. sinica*, only two inversions have evolved that differentiate *C. chloris* from *C. sinica* – neither of which occurred following the divergence between *C. ambigua* and *C. chloris*. *I have treated the three members of a black-headed greenfinch species complex (*C. ambigua, C. monguilloti*, and *C. spinoides*) as a single species *C. ambigua* here based on the lack of any observed premating isolation where their ranges overlap [51].

### Genomic Distribution of Chromosome Inversions

The fastest rearranging autosomal chromosome evolved 4× faster than the slowest (Tables 4, S8). The three top models to explain the variation in inversion fixation rate between the autosomes had nearly equivocal AICc scores and model weights (ΔAICc <2; S10 Table). I used model averaging to combine them into a final model that included only chromosome GC content and repeat density (S10 Table). Of these, only repeat density was found to be significant (z value = 2.3, p = 0.02; S10 Table). The significance of the observed association was not robust to model averaging the six top models with ΔAICc <4, suggesting only a weak effect of repeat density (S10 Table). Inclusion of the Z chromosome in these analyses further reduced the fit of any mutagenic model to explain variation in inversion fixation rate between chromosomes.

**Table 4.** Genomic distribution of pericentric inversions. Autosomes are listed in order of descending size with their presumed homology to the collared flycatcher (*Ficedula albicollis*) genome given in parentheses. Values for chromosome size and map length come from the collared flycatcher genome [44] while GC content and repeat density come from the zebra finch (*Taeniopygia guttata*) genome [55,56]. Variance in branch lengths by chromosome reflects species with missing data.

| Chromosome | Size (Mb) | Map Length (cM) | GC Content (%) | Repeat Density (%) | Branch Length (My) | Inversions |
| --- | --- | --- | --- | --- | --- | --- |
| 1 (FAL2) | 157.4 | 320 | 39 | 0.38 | 4449.2 | 39 |
| 2 (FAL1) | 119.8 | 245 | 39.2 | 0.23 | 4449.2 | 71 |
| 3 (FAL3) | 115.7 | 230 | 39.4 | 0.38 | 4449.2 | 80 |
| 4 (FAL1A) | 74.8 | 230 | 39.7 | 0.14 | 4449.2 | 108 |
| 5 (FAL4) | 70.3 | 175 | 39.2 | 0.28 | 4437.3 | 104 |
| 6 (FAL5) | 64.6 | 172 | 40.8 | 0.12 | 4398.1 | 79 |
| 7 (FAL7) | 39.3 | 125 | 41.1 | 0.14 | 4242.1 | 48 |
| 8 (FAL6) | 37.2 | 122 | 41.6 | 0.19 | 4088.8 | 35 |
| 9 (FAL8) | 32 | 95 | 41.3 | 0.5 | 3973.8 | 23 |
| Z | 59.7* | 165 | 39.2 | 1.5 | 4449.2 | 121 |
| W | 27 |  | - |  | 3387.8 | 100 |

\*The sizes of chromosomes are nearly identical between the collared flycatcher and the zebra finch except for chromosome Z (59.7Mb vs. 74.6Mb, respectively).

Compared to the autosomes, inversion fixation occurred on average 1.5× faster on the Z chromosome and 2.5× faster on the W chromosome (two-sample paired *t*-tests with the 80 clades as replicates, ChrZ: t_79_ = 2.2, p = 0.034; ChrW: t_74_ = 4.0, p = 1.0 × 10^−4^). The W chromosome carried more pericentric inversions than the Z (paired *t*-test: t_74_ = 2.5, p = 0.013). When scaled by chromosome length, the difference on the W chromosome is more dramatic with 3.7 inversions per Mb compared to 1.1 inversions per Mb on average for all other chromosomes, including the Z. The genomic distribution of segregating inversion polymorphisms within species, as well as variants apparently fixed in different populations of the same species, was also biased towards the sex chromosomes. Of the 43 within-species pericentric inversion variants identified from the cytological data, 14% were on the Z chromosome (6 of 43 variants) and 26% on the W chromosome (11 of 43; S2 Table).

## Discussion

Large pericentric inversions have evolved often in passerine birds, with a minimum estimate of one inversion for every 5.3 million years of evolution. This estimate does not account for possible back-substitutions. Within 80 passerine clades, which include more recent events only and so the influence of structural back mutations is minimized, the average rate of pericentric inversion fixation for all chromosomes was one inversion fixed every 3.9 million years, which approximates the rate at which measurable hybrid infertility in birds appears [57], but ranged from no inversions fixed over the span of 27.7My (in the family Dicruridae) to one inversion fixed over 1.2My (in the genus *Chloris*; S5 Table). Many inversions have doubtless gone undetected. Twice as many individuals were karyotyped in the 31 passerine species found to have inversions segregating versus the study as a whole (9.9 versus 4.8 individuals, respectively; excluding 3 species of large sample size, Fig. 5; S2 Table). Moreover, as I have restricted my analysis to only those pericentric inversions large enough to be detectable via cytological analysis (i.e. excluding small pericentric inversions as well as all paracentrics) these counts are surely an underestimate of the true extent of chromosome inversion variation in passerines, as has become clear from genomic studies [44,45,58].

**Fig 5.**
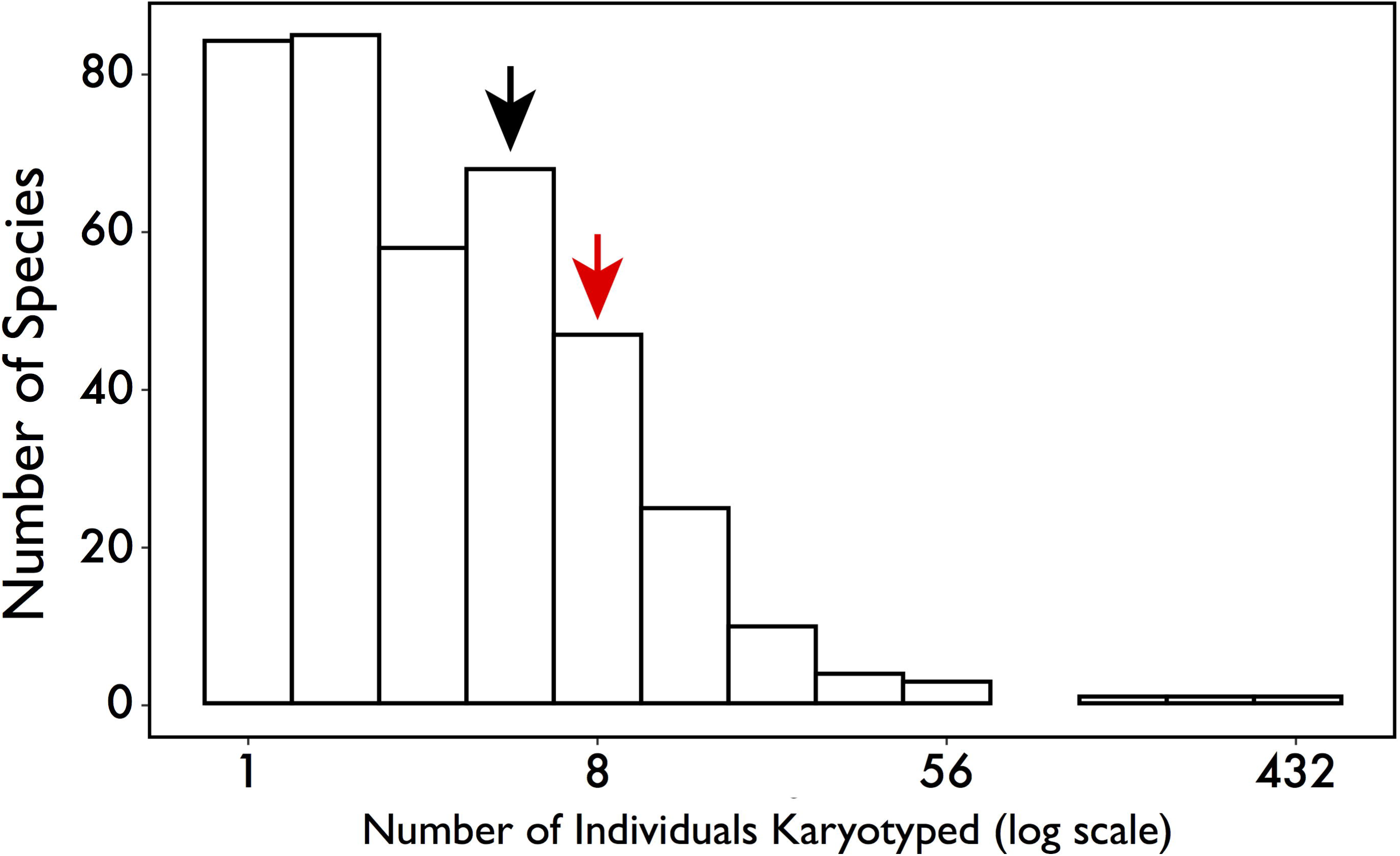
Natural log distribution of sample size for karyotyped species. The average number of individuals karyotyped are indicated by arrows for all species (black arrow, 4.8 indiv.) and species where structural variants were observed (red arrow, 9.9 indiv.) after removing the three species where sampling effort was designed to study inversion polymorphism [S1-2 Tables].

The main finding from this study is that the strongest correlate of inversion fixation after accounting for time, is not range size, but range overlap. Variation in the number of inversions observed across clades positively scales with the proportion of sympatric species pairs each contains and sister species comparisons directly invoke range overlap as an important correlate. Indeed, the evidence suggests that while rearrangements occur more often in clades with larger ranges, thereby rejecting models based on drift and peak shifts [20,21,25], this effect is secondary to whether or not the ranges of these species are sympatric. Range overlap is pertinent to one particular model of inversion spread in which gene flow between partially reproductively isolated forms favors inversions, because F1 hybrids more rarely recombine parental allelic combinations in the inverted region compared to collinear chromosomes. Altogether, the evidence suggests that pericentric inversions in passerines are adaptive and derive their selective advantage by keeping sets of locally adapted alleles together when gene flow between incipient species would otherwise break them up. I first consider caveats before returning to the main results.

The first issue is whether sympatry generally reflects a historical capacity for gene flow that is precluded by allopatry. While species with allopatric ranges may well have been largely allopatric since they first split, species with sympatric distributions may have become reproductively isolated before establishing secondary contact. Empirical evidence across a broad spectrum of taxonomic groups, however, suggests otherwise. Secondary contact between incipient species has regularly been followed by hybridization and genetic exchange (reviewed in [59]). We know for birds that complete postmating isolation in the wild typically requires 3My or more [37,60], with hybrid zones regularly forming between taxa separated by that age ([60], S6 Table). This suggests that secondary contact often precedes the completion of reproductive isolation and could often select for chromosome inversion. Non-hybridizing sympatric pairs exhibit no discernible disparity in the extent of inversion differentiation when compared to hybridizing pairs (Fig. 3; S6 Table).

The second issue is that allopatric sister species may exhibit reduced inversion differentiation due to a lower mutational input because they are younger–less time for an inversion mutation to occur–or because they have smaller population sizes-lower inversion mutation rate per generation. However, I find no strong evidence that the phenomenon of reduced inversion differentiation among allopatric species is primarily a result of allopatric sister species tending to be younger and smaller in range than their sympatric counterparts (Table 3). Analysis of species triplets, which entirely control for age, also find no evidence that the extent of inversion differentiation between sister species and an outgroup taxon is a result of differences in sister range size (S7 Table). Together with the findings from hybridizing sister pairs, these results strongly suggest that range overlap makes an important contribution to chromosome inversion fixation.

### Evaluating Support for Alternative Models of Inversion Evolution

Four alternative models of inversion fixation were considered to explain the distribution of pericentric inversions in passerines. First, in agreement with an earlier analysis [25], I find that genetic drift is unlikely to have been a strong force. The fixation rate for inversions on all chromosome classes across the 80 passerine clades examined scales with body size corrected range size (autosomes: r = 0.38, chromosome Z: r = 0.32, and chromosome W: r = 0.25; S5 Table) while fixation rate by drift should, to a first approximation, be population-size independent [20–22,25].

Assuming rates of evolution by breakpoint selection and meiotic drive models should be largely dependent on mutagenic input, results do not support these models either, because range size is not a strong correlate, and because of the distribution across chromosomes (see below). Indeed, no mutagenic expectation of inversion fixation well explained the genomic distribution of autosomal pericentric inversions across the 410 passerine species examined in this study (Tables 4, S10). These findings suggest that a force beyond raw mutagenic input is responsible for heterogeneity in the extent of pericentric inversion differentiation observed in passerines. Indeed, results are consistent instead with an adaptive model in which an inversion has a selective advantage if it maintains - through recombination suppression - the a) ecological or b) reproductive differentiation of a population at risk of being homogenized by gene flow.

### Gene Flow and Chromosome Inversion Fixation

The adaptive model of rearrangement evolution presented by Kirkpatrick and Barton [31] relies on gene flow between ecologically differentiated populations to facilitate the spread of an inversion. Under this model, an inversion that encompasses two or more loci carrying alleles favored in the environmental background of a population may be favored if recombination would otherwise break them up. One prediction is that species prone to ecological speciation may be more likely to fix inversions first, because divergent selection on loci involved in local adaption is strong and second because diverging forms are more likely to occur in sympatry, being ecologically differentiated, before postmating isolation is complete [61]. For example, finch-like forms appear to speciate ecologically and achieve sympatry more quickly than insectivores [61], so the model of local ecological adaptation predicts finches should carry more inversion differences. While classifications of ecology in this study were crude, feeding guild and inversion accumulation are not correlated (S7 Table). Future empirical efforts to examine the genic content and targets of selection within inversions, like those in flies [62,63], mosquitoes [64], butterflies [6,12,65], sticklebacks [13,66], and monkeyflowers [11,18] will be more powerful ways to assess the degree to which ecology has shaped the evolutionary trajectory of inversions in passerines.

In an extension of the local adaption model, hybridization between incipient species creates a selective advantage for a chromosome inversion that maintains linkage between loci locally adapted to the genetic backgrounds of hybridizing taxa. For example, alleles at loci involved in pre-and/or postmating isolation are favored when kept in tight linkage by an inversion because of the increased production of unfit hybrids when they recombine. The likelihood of inversion differentiation under this model is expected to be most strongly associated with the frequency with which gene flow is part of the speciation process. As geographic isolation precludes hybridization, there is no selective advantage for a novel inversion to capitalize on and species that achieve reproductive isolation in allopatry should maintain chromosome co-linearity. This model has support from both clade and sister species analyses.

Two extreme examples illustrate the case for a role of range overlap in inversion fixation. First, tits (family Paridae) in the genera *Periparus* and *Pardaliparus* last shared a common ancestor 7Ma (5.2 – 8.9Ma, 95% HPD), have largely allopatric distributions (no pair of species overlap in range more than 20%), and no known inversion differences. In stark contrast, an Asian clade of tits in the genus *Poecile* diverged 4.3Ma (3.1 – 5.6Ma, 95% HPD), are largely sympatric (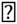 of pairs), and the species examined differ by up to seven pericentric inversions (S5 Table). Natural hybrids between *Poecile montanus* and *P. palustris* have been recorded in the wild [51], more directly linking gene flow to inversion evolution. A second example comes from greenfinches in the genus *Chloris* (family Fringillidae, Fig. 4). Inversion differentiation between *C. sinica* and sympatric *C. ambigua* has outpaced inversion differentiation between *C. sinica* and allopatric *C. chloris* (Fig. 4). Further suggesting a positive interaction between gene flow and inversion evolution, *C. ambigua* and *C. sinica* hybridize at low frequencies where their ranges overlap [51].

### Genomic Distribution of Pericentric Inversions

The distribution of chromosome inversions detected using comparative genomic approaches in birds is positively associated with chromosome size [44,45], and inversion breakpoints are often located in regions with elevated recombination rates, GC content, and repeat density [44]. These results were not replicated here. A primary reason for this difference likely resides in the different size classes of inversions considered between studies [25]. Inversions detected from comparing high-resolution linkage maps [44] or whole genome alignments [45] are capable of finding structural variants orders of magnitude smaller than the exclusively large inversions I identified from cytological data. Indeed, the pericentric inversions considered here may be of greater evolutionary relevance as they potentially come with both higher fitness costs and greater selective advantages than the more comprehensive set of inversions found in comparative genomic surveys.

However, sex chromosomes do accumulate pericentric inversions more rapidly than the autosomes (Tables 4, S8), despite having smaller population sizes. A quarter of all identified inversion polymorphisms occur on the W chromosome while 14% are on the Z chromosome. The higher rate of sex chromosome inversion evolution could be considered a consequence of sex chromosomes possessing a higher structural mutation rate; lower fitness costs for structural rearrangements, a greater influence of genetic drift, or - in the case of the Z chromosome - a greater fitness benefit to a recombination modifier. I find no evidence for an influence of drift in governing variation in inversion fixation on any chromosome or for a higher mutation rate on the Z compared to the autosomes. Inference about the W chromosome is difficult as its features are not well understood. In contrast, a recent study finds that positive selection is the driving force responsible for elevated rates of functional differentiation observed for Z-linked genes in six species of Galloanserae [67]. These are ideal conditions for inversions on the Z chromosome to be favored as recombination modifiers when differentiation occurs with gene flow. One route to further understanding will come from explicit studies of the selective forces maintaining inversion polymorphisms within species [7–9].

## Materials and Methods

### Identifying Inversions

I called chromosome inversions from classic studies of gross karyotype structure that encompass nearly 8% of all passerine species and >50% of passerine families. Of the 427 passerine species that have had their karyotypes described, I discarded 15 because the cytological data was not of sufficiently high quality to include in this study and two because no suitable genetic data currently exists for them and no tissue materials were available. I analyzed cytological data for the remaining 410 species, representing birds from 59 families (S1 Table). Data was sourced from 110 studies that span five decades of cytological research (S1 Table). Methods utilized to describe karyotype varied from simple Giemsa staining to fluorescent *in situ* hybridization with chromosome painting. Sampling rigor varied across studies with respect to the number (average of 7 karyotyped individuals per taxon, range from 1 to 432; Fig. 5) and sex representation of each species (data from both males and females in 296 of 410 species; S1 Table). Sampling information was not given for 29 species (S1 Table). Due to the considerable heterogeneity in the quality and quantity of karyotype descriptions between species and studies, I focus on a simple yet powerful trait with which to infer chromosome inversion differences between and within species: centromere position.

For each species, I converted centromere position for the 9 largest autosomal chromosomes and both sex chromosomes into character state data (S1 Table). I scored each chromosome for approximate centromere position (i.e., whether they were metacentric, sub-metacentric, sub-telocentric, or telocentric), following conventions established by Levan et al. [68]. I identified homologous chromosomes between species by a combination of their physical size, shared banding pattern, and matching chromosome painting, as the information was available. I treated centromere position of a chromosome as distinct when species shared the same general classification (e.g., both were sub-metacentric) but the authors noted that the banding pattern flanking the centromere consistently differed. I only include pericentric inversions in my analyses as the cytological data has far less power to identify paracentric inversions (those not encompassing the centromere). Centromere repositioning can result from processes other than pericentric inversion, such as the redistribution of heterochromatin [69,70] and the evolution of neo-centromeres [70,71]. I found no evidence, however, for either of these alternative mechanisms of centromere repositioning in the 85 species with banding data available to test for them as centromere movement was supported by inversion of proximal banding patterns (Table S1).

While the distribution of fixed inversion differences can be used to infer historical patterns of selection, the mechanisms of selection affecting inversions are best studied when rearrangements still segregate in natural populations. I therefore evaluated all species for the occurrence of pericentric inversion polymorphisms and for the presence of inversions present in different parts of species ranges (S1-2 Tables). Polymorphisms segregating within populations were often noted in the paper of interest, but the majority of geographic variants are first reported in this study, as they generally depend on comparing different published studies (S2 Table). Of the 50 total rearrangement polymorphisms identified, two are likely a product of chromosome translocation and three are shared between species - two across three species and one between two species (S2 Table).

### Phylogenetic Analyses

In order to characterize the phylogenetic distribution of chromosome inversion fixation, I built a time-dated phylogeny for the 410 passerine species with karyotype data available. I gathered sequence data from six genes: two mitochondrial: *cytb* and *ND2*, and four nuclear: myoglobin (*MG*) exons 2–3, ornithine decarboxylase (*ODC*) exons 6–8, beta-fibrinogen (*FIB5*) exons 5–6, and recombination activating protein-1 (*RAG1*). Data were primarily sourced from GenBank. For 12 karyotyped species with no or low sequence representation I generated the data myself using standard methods (Table S3). Phylogenetic and dating analyses were conducted using BEAST v1.8.2 [72]. Sequence data was partitioned by locus, each with its own uncorrelated lognormal relaxed clock, and assigned the optimal-fit model of sequence evolution estimated for each locus using jModelTest v0.1.1 [73]. The phylogeny was time-calibrated using 20 fossil calibrations broadly dispersed both in time and topology (S1 Fig.; S4 Table). This is, to my knowledge, the most extensive fossil calibration effort to date within Passeriformes. Each fossil calibration was applied to its corresponding node as a minimum age bound using a conservative uniform prior based on the age of the fossil itself and 80Ma. I ran BEAST for 50 million generations and sampled every 5000 for a total of 10,000 trees of which the first 1000 were discarded as burn in. I assessed run length and appropriate sampling for each parameter using Tracer v1.6 [74]. Using TreeAnnotator v1.7.2 [72], I extracted the maximum clade credibility tree, with associated confidence intervals for median node heights (Figs. 2, S2).

### Phylogenetic Distribution of Inversion Fixation

In order to map inversion evolution across the phylogeny, I estimated the ancestral centromere position (up to 4 possible states: metacentric, sub-metacentric, sub-telocentric, or telocentric) for each chromosome at each node in the tree by maximum likelihood in Mesquite v2.7.5 [75]. I obtained the maximum likelihood estimate for each ancestral centromere position for each chromosome at every node. Inversions were inferred to have occurred upon branches where the karyotype of an internal node differed from subsequent nodes or the tips and was supported by a maximum likelihood, p > 0.75. I used this phylogenetic representation of inversion evolution in passerines to investigate the drivers of inversion fixation between species and within the genome. I conducted analyses at two different phylogenetic levels. First I defined 80 clades comprising between 3 to 85 species and, second, I used sister species pairs.

### Chromosome Inversion Variation Between Clades

I partitioned the phylogeny of karyotyped taxa into 80 clades of closely related species in order to examine the factors associated with broad scale variation in chromosome inversion evolution. Many clades contain additional species that were not karyotyped, and hence not included in the tree, yet these species may influence chromosomal evolution in the focal taxa, e.g. through range overlap. To take this into account, I utilized phylogenies from 54 published family-level studies in order to determine which non-karyotyped species to include in clade level analyses (S5 Table). Clades were assigned based on the following grouping criteria: the two most distantly related karyotyped species were less than 15 million years diverged, member species’ were the result of speciation within a single geographic region (i.e. all clade members speciated in Australia), member species were ecologically similar (i.e. finches, warblers, frugivores, nectarivores, or omnivores), a comprehensive family level phylogeny exists to identify non-karyotyped member taxa, and they encompassed at least three species including non-karyotyped taxa. After filtering based on the above criteria, 285 of 410 karyotyped species were assigned to 80 clades (S1, S5 Tables).

I measured variation in karyotype evolution across passerine clades by counting the total number of inversions that had fixed on each chromosome, summing over all branches within the clade. I did not include inversion polymorphisms in this count unless the ancestral conformation of the chromosome polymorphic for an inversion, determined in Mesquite, was neither of the segregating forms. I calculated clade branch length as the sum of branch lengths for species with centromere position scored at each of the 9 autosomes, the Z, and W chromosomes. For example, if all species within a clade had complete karyotype records (i.e. centromere position scored for all 11 chromosomes), the branch length value of that clade was the sum of all branches multiplied by a factor of 11. For species missing data for a chromosome, the length of the branch leading to that species was removed from the clade total according to the total number of missing chromosomes (i.e. if a species was missing data at two chromosomes then 2× the branch length to that species were subtracted from the clade total).

I collected range overlap, range size, and body mass data from the complete taxon set for each clade (i.e. including both karyotyped and non-karyotyped species) in order to evaluate the extent to which variation in demography (population size) and speciation history (range overlap) has impacted inversion evolution (S5 Table). I extracted range data for all species from natureserve.org using the programs Sp [76] and PBSmapping [77] in R. I assigned each clade a range size value corresponding to the median range size (km^2^) of all member taxa. Median body mass (g) for each clade was calculated from Dunning [78]. I used range size together with body mass in mixed models as proxies for population size based on the positive relationship between the geographic area a species occupies and its nucleotide diversity [79–82] and the negative relationship generally observed between body size and population density [83]. I assigned a range overlap score to each clade based on the proportion of all species pairs whose ranges overlap by >20% and/or are known to hybridize in the wild [51]. I include hybridizing taxa together with taxa whose ranges are sympatric because both imply there is at least the potential for gene flow between taxa. Lastly, I considered a broad role for ecology on chromosome inversion evolution across clades according to the feeding guild used when defining clades (i.e. clades defined as comprising granivores, insectivores, frugivores, or ominvores; [33]).

In order to improve the interpretability of regression coefficients, the total number of inversions, branch length, range size, and body mass were log transformed, range overlap was arcsine square root transformed, and all variables were centered before analysis [84]. I then evaluated the extent to which the number of inversions that had fixed in each clade was associated with branch length, range overlap, range size, body mass, and ecology using generalized least squares to take into account phylogenetic relationships [85]. To do this, I used the NLME package in R [86], with the expected error covariance matrix computed based on the phylogenetic distances between clades (S3 Fig.). To assess the relative importance of each factor on the number of inversions fixed in each clade, I compared all possible models and selected the best-fit model based on sample size-corrected information criteria (AICc) using the R package MuMIn [50].

### Chromosome Inversion Variation Between Sister Species

I also considered the distribution of inversions between sister species, including both fixed differences and inversions segregating in one taxon but not the other. I determined which karyotyped species pairs were true sisters using the available phylogenetic literature (S6 Table). I considered a sister pair to hybridize if they had documented hybrid zones or extensive natural hybridization where they co-occur [51]. In total, I identified 47 true sisters with both species karyotyped, of which 12 are known to regularly hybridize in nature (S6 Table).

For all 47 sister pairs, I calculated the number of inversion differences between them, their time to common ancestry, average range size, range overlap, and whether they are known to hybridize in the wild. Inversion differentiation was scored both as a binary character (no inversions or at least one inversion difference) and as a count (total number of inversion differences). Range overlap was evaluated as a binary character: no overlap or some overlap. I only used this binary categorization because subdividing sisters who overlapped in range into either parapatric (< 20% overlap) or sympatric (>20% overlap) bins did not improve the fit of any model or alter the results in any way. I used a linear model to examine the interaction between the number of sister pair inversion differences and each factor (age, range size, range overlap, and hybridization) after transforming the continuous character data as described for analysis of clades. Lastly, I assessed whether sister species with overlapping ranges, and the subset of sympatric sisters known to hybridize, are more likely to differ by an inversion than allopatric sisters using *t*-tests.

Genetic distance is not time but rather an estimate of time, and one that can come with substantial error. This error can diminish the true contribution of time and elevate the importance of alternative factors [52]. A method to completely control for the potentially confounding influence of time is the use of species triplets [52–54]. A triplet consists of a sister species pair (A, B) and a single outgroup taxon (O). Both sister taxa have by definition been separated from the outgroup for the same length of time. If O overlaps B but not A, then the presence of more inversion differences between O and B than O and A gives strong support for a role of range overlap independent of time (see Fig. 4). I assembled a set of species triplets from the phylogeny of karyotyped species and published phylogenies, using the following criteria: both sister species A and B have been karyotyped, A and B are allopatric, and B overlaps in range with O but species A does not. This resulted in just 5 triplets (S7 Table). I relaxed the criteria to allow 1) range overlap between A and B and 2) range overlap between A and O so long as they were not sympatric (i.e. ranges overlapped less than 20%) and overlapped in range less than B and O. The average extent of range overlap between species A and O, when they did overlap, was 3× less than the extent of range overlap between B and O (S7 Table). Nineteen triplets were present after applying the relaxed filtering criteria (S7 Table).

I counted the number of inversions inferred to have occurred along the branches leading to species A and B, respectively, based on the distribution of fixed inversions in the complete karyotyped species phylogeny. I also included inversion polymorphisms found in one but not the other taxon. I scored each triplet as follows: more inversions in A than B, more inversions in B than A, or no difference in the number of inversions between A and B. I evaluated the direction and significance of the relationship between range overlap and inversion evolution across all triplets by applying a signed rank test to those sisters where the number of inversions differed.

### Genomic Distribution of Chromosome Inversions

Inversion fixation models that depend heavily on mutational input (e.g. meiotic drive and breakpoint selection) predict a strong correlation with range size but they also predict a strong association with mutation rate. In a final analysis to examine the extent to which inversion evolution is a mutation limited process, I examined the distribution of chromosome inversions across the genome and evaluated the degree to which the number of inversions fixed on a chromosome (S8 Table) was associated with four possible mutagenic processes. First, if the mutation rate for inversions is constant per DNA base, the number of inversions should be proportional to chromosome size. Second, because inversions are derived from double-stranded meiotic breaks, the number of inversions on a chromosome could best be predicted by its map length or GC content - features associated with the number of cross-overs per chromosome [87,88]. Third, as inversion breakpoints are often located in repeat-rich regions of chromosomes [43,44], I tested for an association between the number of inversions and a chromosome’s repeat density. Fourth, I asked if the dynamics of inversion fixation on the sex chromosomes and the autosomes differ [25]. Mutation rates on the Z chromosome should be relatively high in birds because the Z spends 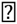 of the time in males, however this mutational advantage needs to overcome the fact that there are only ¾ as many copies of the Z as each of the autosomes [89,90]. In contrast, the W chromosome should have a low mutation rate both because it spends all its time in females and there are ¼ as many copies of the Was the autosomes.

Primary estimates of chromosome physical size, map length, and GC content were derived from the collared flycatcher genome assembly and linkage map [44] and chromosome repeat density was estimated from a RepeatMasker annotation of the zebra finch genome (http://www.repeatmasker.org; [55]). I use chromosome size and map length data from the collared flycatcher but obtained identical results when analyses were repeated using chromosome size and map distance data derived from zebra finch [42,56] and hooded crow (Corvus cornix [91]; S10 Table). Comparative genomic studies indicate that chromosome size (excluding the W chromosome), GC content, and repeat density are conserved even between species in different avian orders [44,45,92]. While the recombination landscape may have phylogenetic signal [93–95], recombination hotspots are well maintained in passerines [58].

I used data from all 410 karyotyped species to examine the correlation between a chromosome’s inversion fixation rate and its physical size, GC content, repeat density, and map length, using each chromosome as a replicate. In order to account for species with missing data I use inversion fixation rate (total number of inversions fixed on a chromosome divided by the combined branch length for all species with data for that chromosome) rather than inversion number (S1 Table). Independent variables were log-transformed. I evaluated support for alternative mutagenic hypotheses by comparing between all possible linear models and selected the best-fit model using the R package MuMIn [50]. Restricting the analysis to the 291 species with complete karyotype data (i.e. documented centromere position for all 11 chromosomes) yielded a similar result (S10 Table). Finally, I tested for significant differences in the rate of inversion fixation between the autosomes, Z, and W chromosomes using the 80 independent passerine clades defined above as replicates and paired *t*-tests.

## Acknowledgements

I thank N.S. Bulatova, B.S.W. Chang, E.J. de Lucca, G. Semenov, P. Tang, I.M. Ventura, Y. Wu, and the University of Chicago Library for their help in accessing cytological studies not currently available online. I thank S.G. Dubay, T.D. Price, Supriya, and A.E. White for their assistance with statistical analyses and figure aesthetics. Tissue materials for species without data on GenBank came from the Kansas University Biodiversity Institute and Natural History Museum. Thanks to M. Sorenson and C. H. Oliveros for sharing unpublished phylogenetic results. A.E. Johnson provided original artwork of the Chloris greenfinches used in Fig. 2. I am grateful to the assistance of T.D. Price for his helpful comments and suggestions on multiple versions of this manuscript and to M. Przeworski for her feedback on a single version.

## Supporting Information

**S1 Fig. Phylogenetic distribution of fossil calibrations.** Calibration nodes numbered following S4 Table and labeled (red circles). Species included solely for calibration purposes colored red.

**S2 Fig. Pericentric inversion fixation rate variation across passerine birds.** The phylogenetic relationships between the 410 karyotyped species in this study are presented in a time-dated maximum clade credibility tree. Branches are color-coded according to the inferred rate of pericentric inversion fixation using the R package ggtree[48]. Rates are partitioned according to the Jenks natural breaks method where variance within bins is minimized, while variance between bins is maximized [49].

**S3 Fig. Phylogeny used in phylogenetic generalized least squares for clade level analysis.** Branch lengths are proportional to time and were used to compute the expected error covariance matrix between clades for phylogenetic generalized least-squares. Species identity for the 285 taxa assigned to clades is given in S1 Table. Clade information (inversions, branch length, range size, range overlap, ecology, etc.) are given in S4 Table.

**S1 Table. Chromosome character state matrix with references.**

**S2 Table. Chromosome rearrangement polymorphisms.**

**S3 Table. Loci used in phylogenetic analyses.**

**S4 Table. Fossil calibration set with references.**

**S5 Table. Clade data.**

**S6 Table. Sister species data.**

**S7 Table. Species triplet data.**

**S8 Table. Genomic distribution of inversions across 80 passerine clades.**

**S9 Table. Model comparison results for clade and sister species analyses.**

**S10 Table. Model comparison results for genomic distribution of inversions.**

